# Human-level saccade detection performance using deep neural networks

**DOI:** 10.1101/359018

**Authors:** Marie E. Bellet, Joachim Bellet, Hendrikje Nienborg, Ziad M. Hafed, Philipp Berens

## Abstract

Saccades are ballistic eye movements that rapidly shift gaze from one location of visual space to another. Detecting saccades in eye movement recordings is important not only for studying the neural mechanisms underlying sensory, motor, and cognitive processes, but also as a clinical and diagnostic tool. However, automatically detecting saccades can be difficult, particularly when such saccades are generated in coordination with other tracking eye movements, like smooth pursuits, or when the saccade amplitude is close to eye tracker noise levels, like with microsaccades. In such cases, labeling by human experts is required, but this is a tedious task prone to variability and error. We developed a convolutional neural network (CNN) to automatically detect saccades at human-level performance accuracy. Our algorithm surpasses state of the art according to common performance metrics, and will facilitate studies of neurophysiological processes underlying saccade generation and visual processing.

## Introduction

Eye tracking is widely used in both animals and humans to study the mechanisms underlying perception, cognition, and action, and it is useful for investigating neurological and neurodegenerative diseases in human patients^1–5^. This is in part due to practical reasons: recording eye movements is relatively easy^6^, while, at the same time, eye movements can be highly informative about brain state^7, 8^.

The most prominent type of eye movement, in terms of eyeball rotation speed, is a ballistic shift in gaze position, called saccade. This type of eye movement occurs 3–5 times per second, and it can realign the fovea with interesting scene locations within only ∼50 ms. Naturally, saccades cause dramatic changes in visual input when they occur, and they therefore impact neural processing in different visual areas and also in a variety of ways^9–17^. This even happens for the tiniest of saccades, called microsaccades, that occur when gaze is fixed^18–27^. Therefore, studies not quantitatively analyzing microsaccades can miss important behavioral and neural modulations in experiments^28^. Saccades and microsaccades are, additionally, key discrete events in eye tracking traces that can be useful for parsing other eye movement epochs (e.g. smooth pursuits, ocular drifts, ocular tremors) for further analysis. Therefore, detecting saccades is typically the first step in any quantitative analysis of behavior or neural activity that might be impacted by these eye movements.

Several algorithms have been proposed for automating the task of saccade detection (reviewed by ^29^). For example, Engbert and Mergenthaler developed a method for classifying sac-cades and microsaccades based on an adaptive threshold^30^. This algorithm (which we refer to here as EM) is particularly popular because of its simple implementation and ease of use, as well as its ability to detect even microsaccades. However, this algorithm, like others, may still mislabel some microsaccades due to high eye tracker noise (as is typical with video-based eye trackers) as well as small catch-up saccades occurring during smooth pursuit. Other existing algorithms^31, 32^ have the added advantage of providing additional labels for fixations and post-saccadic oscillations (PSO).

Despite their success, several shortcomings still render the use of existing algorithms either less reliable than desired or, at the very least, cumbersome. While the performance of many published algorithms is promising^29, 32^, it does not reach the level of trained human experts. Also, none of the existing algorithms shows convincing performance for all eye movement-related events that may need to be analyzed (e.g. fixations, saccades, PSO, blinks, smooth pursuits). In addition, equipment-dependent hyperparameters, such as thresholds, need to be chosen for most algorithms, a fact that renders broad usability difficult. For example, even simple changes in eye tracking hardware, involving changes in sampling frequency or measurement noise, require re-tuning of such parameters. Re-tuning is also needed when the ranges of eye movement amplitudes being studied are modified (e.g. microsaccades versus larger saccades). Perhaps most importantly, objective parameter estimation in existing algorithms is currently a challenging task because of a limited amount of available reliably labeled data. Finally, in many cases, applying available online resources is not straightforward. As a result of all of the above shortcomings, current laboratory practice often still involves experimenters spending substantial amounts of time to carefully relabel parts of their data after automatic saccade detection.

Here we propose a convolutional neural network (CNN) for classifying eye movements. The architecture of the network is inspired by U-Net, which has successfully been used for image segmentation^33^. We evaluated our network (U’n’Eye) on three challenging datasets containing small saccades occurring during fixations or smooth pursuits. On these datasets, U’n’Eye reached the performance level of human experts in labeling saccades and microsaccades, while being much faster. The network also beat state-of-the-art algorithms on a benchmark dataset not just for saccade detection, but also for PSO. It can be trained quickly, even on a standard laptop, and its adaptability to different datasets makes U’n’Eye the novel state-of-the-art eye movement detection algorithm. An open source implementation of U’n’Eye is available online (https://github.com/berenslab/uneye), and an easily accessible web service will be provided. Our labeled datasets will also be freely available upon publication.

## Results

### Design of a convolutional neural network for eye movement classification

We developed a CNN that predicts the state of the eye for each time point of an eye trace. The aim of the network was to segment eye movement recordings (Fig. 1) into epochs containing saccades/microsaccades (orange highlights) versus epochs not containing these eye movements (but see also below for additionally classifying PSO using our network). Our primary goal was to have a network that can seamlessly handle the challenging scenarios of tiny microsaccades during fixation (Fig. 1A), small catch-up saccades embedded in relatively high smooth pursuit eye velocities (Fig. 1B), and microsaccades and saccades occurring in recordings with higher noise levels associated with video-based eye trackers when compared to, say, scleral search coil techniques^34, 35^ (Fig. 1C). We therefore trained and tested the network on three different challenging datasets (see Methods and Table 1), which contain labels for fixations and saccades manually determined by human experts. We also tested our network on artificially generated noisy eye movement traces, in which the ground truth for saccade times was known (Methods) (Fig. 1D). Finally, we compared our network’s performance to different existing algorithms, both on our datasets and also on a publicly available benchmark dataset^31^.

**Figure 1:**
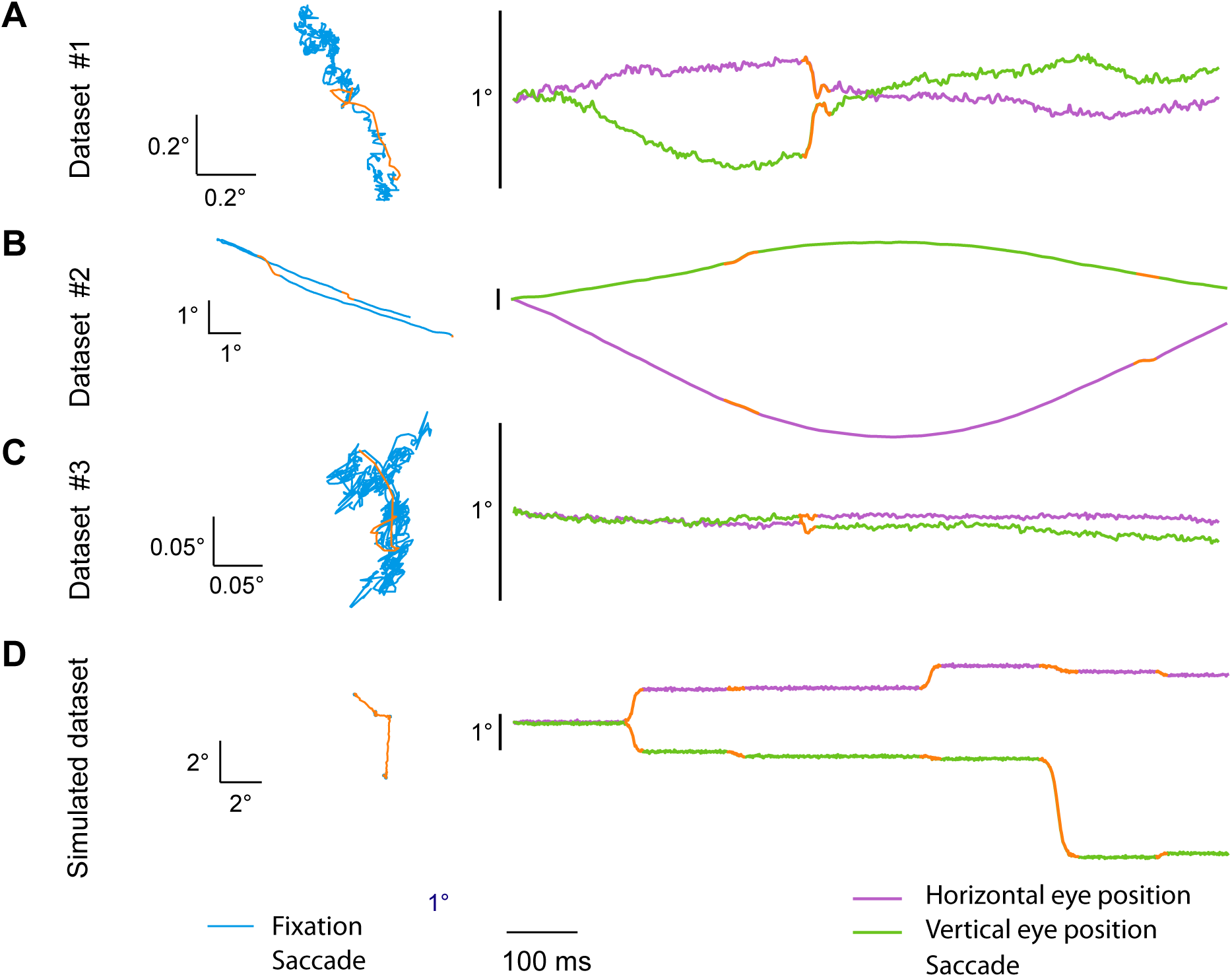
Examples of eye traces containing saccades for detection. (A) Microsaccades during fixation recorded with a video-based eye tracker. (B) Catch-up saccades during smooth tracking recorded with scleral search coils. (C) Microsaccades during fixation recorded with a video-based eye tracker. (D) Simulated saccades. Left panels are the 2D representation of the eye trajectory over 1 second of recording. Right panels are the horizontal and vertical eye position of the corresponding traces on the left, now presented as a function of time; in this case, an upward deflection in the shown traces corresponds to a rightward or upward eye movement for the horizontal and vertical components, respectively. Note that in B, we refer to the non-saccadic smooth change in eye position as “fixation” for simplicity, since the primary goal of our algorithm was to detect saccades, irrespective of whether they happened during fixation or embedded in smooth pursuit eye movements.

**Table 1:** Dataset characteristics. All statistics refer to saccades. Note that minimum saccade amplitude may appear very low due to the existence of some saccades that had very strong dynamic overshoot (a substantial saccadic movement followed by one lobe of a PSO almost to the original eye position before saccade onset).

| Dataset | 1 | 2 | 3 |
| --- | --- | --- | --- |
| Subjects | Humans | Monkeys | Monkeys |
| Eye tracker | Eyelink 1000 | Search coil | Eyelink 1000 |
| Sampling frequency (Hz) | 1000 | 1000 | 500 |
| Saccade type | Microsaccades and memory saccades | Saccades during smooth pursuit | Microsaccades |
| Mean duration ± std (ms) | 44.58 ± 15.42 | 37.51 ± 8.81 | 23.12 ± 6.52 |
| Median duration (ms) | 42 | 36 | 22 |
| Minimum duration (ms) | 11 | 18 | 8 |
| Maximum duration (ms) | 169 | 97 | 54 |
| Mean amplitude ± std ($^{\circ}$) | 0.69 ± 0.93 | 1.07 ± 0.70 | 0.07 ± 0.04 |
| Median amplitude ($^{\circ}$) | 0.43 | 0.96 | 0.06 |
| Minimum amplitude ($^{\circ}$) | 0.02 | 0.04 | 0.003 |
| Maximum amplitude ($^{\circ}$) | 11.34 | 7.03 | 0.38 |
| Mean peak velocity ± std ($^{\circ}$/s) | 102.46 ± 68.82 | 68.23 ± 42.98 | 62.40 ± 19.75 |
| Median peak velocity ($^{\circ}$/s) | 81.91 | 56.59 | 59.57 |
| Minimum peak velocity ($^{\circ}$/s) | 17.81 | 11.49 | 25.50 |
| Maximum peak velocity ($^{\circ}$/s) | 547.72 | 450.44 | 167.77 |
| Median instant. velocity ($^{\circ}$/s) | 5.63 | 15.70 | 12.21 |

The network operates on the eye velocity signal and requires no other preprocessing. Eye velocity is computed as the differential of eye position, and chunks of eye velocity signals are then input to the network. Briefly, the network’s architecture is based on the U-Net, a CNN for pixel-by-pixel image segmentation^33^, which we modified to process one dimensional signals and output a predictive probability for each eye movement class at every time point (Fig. 2A). A major change compared to the original U-Net architecture is that we introduced batch normalization (BatchNorm) layers^36^. BatchNorm layers subtract a mean from their input and divide it by a standard deviation. Both of these parameters are estimated for each layer over mini-batches of training samples during learning. This method normalizes the distribution of activations across the network layers, allowing for higher learning rates and reducing over-fitting (see below)^37^. We also applied a rectified linear unit (ReLu) function between each convolutional and batch normalization layer. The ReLu function, or heavy-side step function, introduces nonlinearities in the network, allowing it to apply arbitrary-shaped functions to the input data. Finally, the U-shaped architecture of the network leads to temporal downsampling and upsampling in the hidden layer representations (Fig. 2). Downsampling is achieved by max pooling (MaxPool) operations that reduce the dimensionality of the network content, extracting relevant features. Upsampling is realized by transposed convolution. Convolutional kernels and max pooling operations together lead to the integration of information over time. Due to the network design, the probability assigned to each time bin can be influenced by *±*89 preceding and following time bins (Fig. 2B). Thus, U’n’Eye takes into account a large enough signal in order to make point predictions of the correct eye movement class.

**Figure 2:**
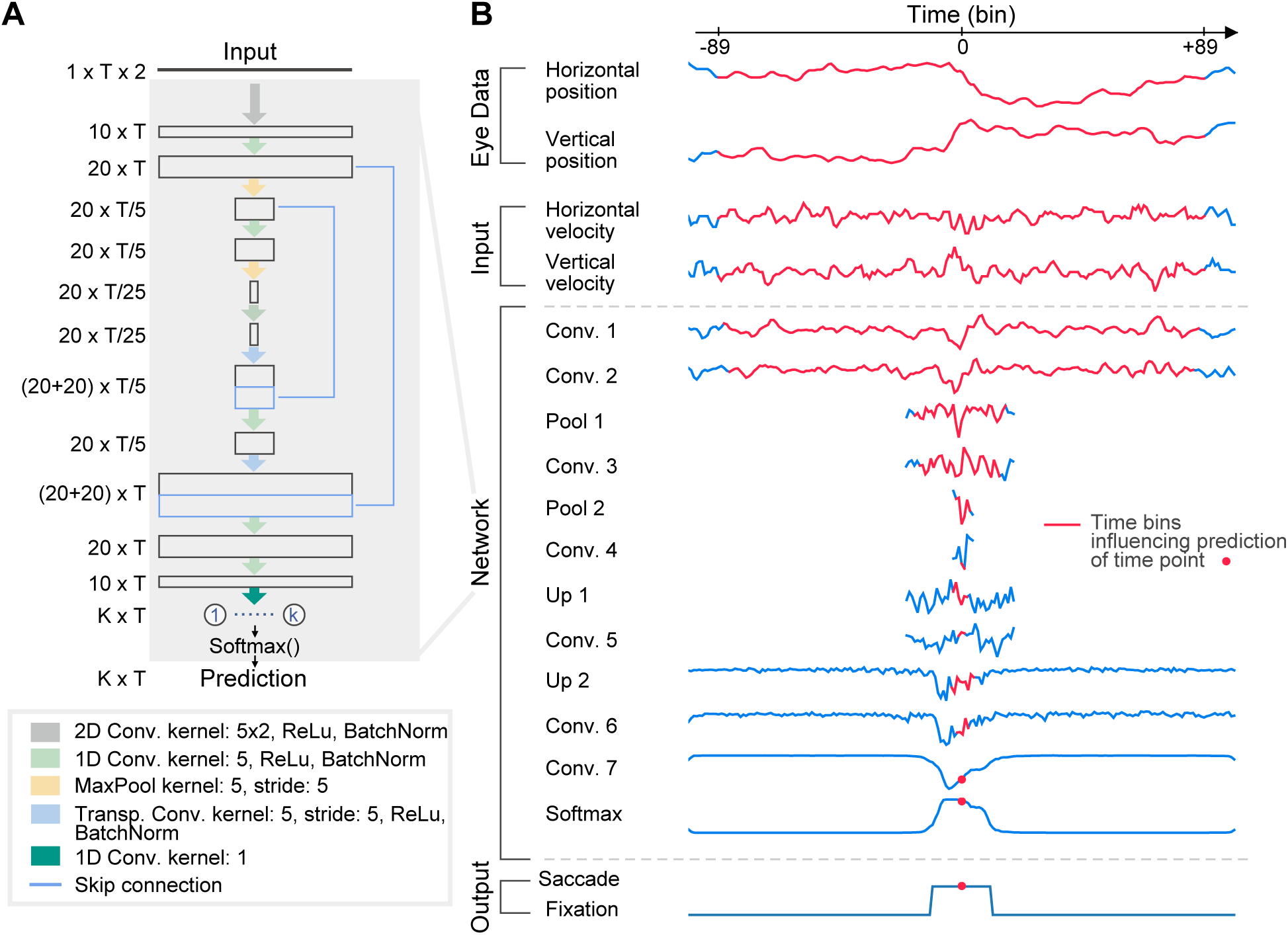
U’n’Eye. (A) Network architecture. The input matrix contains horizontal and vertical eye velocity. T is the number of input time points (see B), and K is the user-defined number of eye movement classes (e.g. “fixation” versus “saccade” in a binary classifier). The different network layers are described in the text. (B) The output probability of one time bin is influenced by 89 time samples before and after this time bin. For each layer of the network, the red color indicates the range of influence of the time bin indicated by the red dot in the output. Traces show the projection of the layer’s output onto its first principal component. The outputs of convolutional layers 6 and 7 resemble the final classifier’s output probability (Softmax), whereas early convolutional layers 1 and 2 seem to perform noise reduction.

### U’n’Eye achieves human-level performance

Our network achieved human-level performance after training on our datasets. We first illustrate this with three example scenarios for detecting saccades (Fig. 3). In the first example, a small microsaccade occurred with substantial oscillation in eye position towards movement end, and with the amplitude of the movement being near the eye tracker noise level (Fig. 3A). The human coder 1 considered the post saccadic oscillation as part of the saccade, and so did our network trained on his training set. On the other hand, coder 2 determined that the saccade ended earlier, and our network trained on his training set did the same. Thus, our network could match the criterion used by an individual human coder very well (Fig. 3A, bottom). Moreover, our network successfully avoided a false detection by the EM algorithm on these traces. In the second example, the EM algorithm missed all three saccades, but our network successfully flagged them (Fig. 3B). Finally, the eye movement in the third example was collected with a video-based eye tracker having substantially more noise (Fig. 3C). In this case, two errors made by the EM algorithm were successfully excluded by our network.

**Figure 3:**
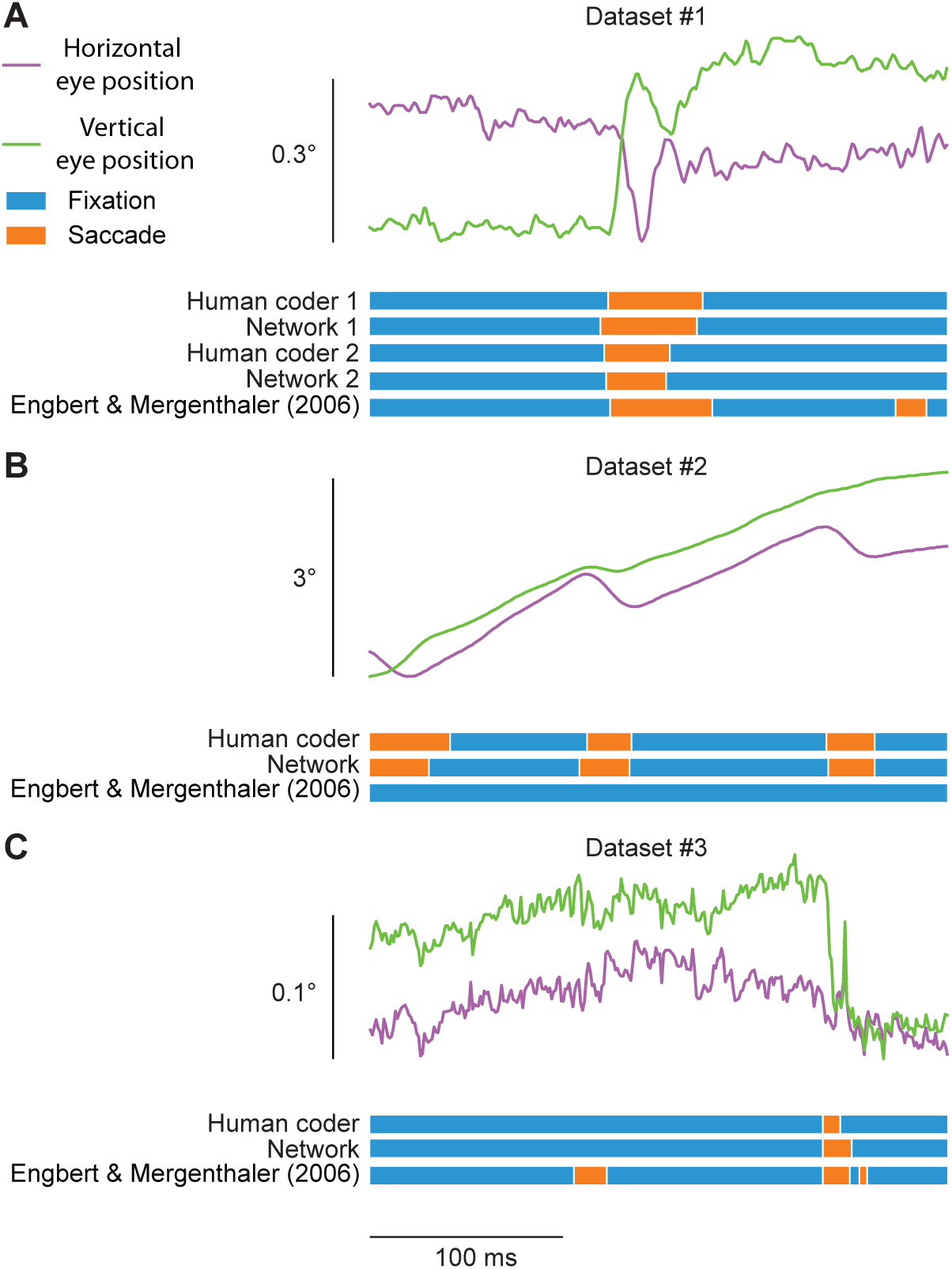
Examples of eye traces from all three datasets, with saccades being labeled by either human coders, different instances of U’n’Eye, or a popular algorithm from the literature. (A) An example microsaccade exhibiting substantial PSO. The top two traces show eye position as a function of time in an identical format to Fig. 1. Below the eye position traces, we show labels for fixation or saccade made by two human experts (coder 1 and 2) as well as predictions of two separate networks. Network 1 was trained on labels from coder 1, and network 2 was trained on labels from coder 2. Note how each network matched the performance of the human coder. The very bottom row shows the performance of the Engbert and Mergenthaler^30^ algorithm (referred to as EM in all remaining figures), which suffered from a false alarm later in the trace due to eye tracker noise. (B) Saccades embedded in smooth pursuit eye movements. Here, our network successfully detected three catch-up saccades, all of which were missed by the EM algorithm. The reason that they were missed is that the saccades were directed opposite to the ongoing pursuit, resulting in momentary reductions in eye speed, as opposed to increases. (C) An example microsaccade embedded in high eye tracker noise. Once again, the EM algorithm suffered from false alarms due to eye tracker noise.

To present more quantitative performance measures, we first tested our network on our in-house datasets. We performed 10-fold cross-validation separately for all three datasets. In each cross-validation round, 90% of the data was used for training the network, and the remaining 10% were used to test performance. A separate validation set from each dataset was used to detect over-fitting of the network. To prevent such over-fitting, we regularized the weights of the network using the L2 penalty^38^ (Methods), preventing the parameters of the network from deviating excessively from zero. Furthermore, we made use of early stopping. For this, a separate validation set was used and the validation set error computed in each epoch. Training was stopped at the point of smallest validation set error. For datasets 1 and 2, 950 sec of eye traces were used for cross-validation and 50 sec for validation. Thus, each training set contained 855 sec of data. For dataset 3, 330 sec were used for cross-validation and 23 sec for validation, resulting in 297 sec of data in each training set. Training took approximately 1 minute per second of training data on a CPU (Methods).

When comparing our network’s performance to the velocity-threshold-based EM algorithm, we again used 10-fold cross-validation to estimate the optimal parameter *λ* that sets the velocity threshold dependent on noise level in that algorithm. As detailed below, we used different metrics to evaluate performance of the algorithms. For the estimation of *λ*, we therefore optimized two different performance metrics which led to slightly different scores of the EM algorithm.

Finally, similarity of the algorithms’ predictions to human labels was evaluated using three metrics. To this end, we calculated Cohen’s kappa, which is a sample-by-sample similarity measure that takes chance agreement of two predictors into account^39^. As a second metric, we calculated the F1 score, which is an accuracy measure that considers precision and recall of a classifier. Recall corresponds to the number of correctly detected saccades divided by the number of saccades that were labeled by the human expert. Precision, on the other hand, is the number of correctly classified saccades divided by the total number of saccades detected by the classifier (Methods). The F1 score is defined as the harmonic mean of both, and it thus only measures how accurate saccades were detected without taking into account their timing (i.e. exact saccade onset and offset times). Correctly labeling saccade onset and offset can be crucial for further analyses. Therefore, we additionally computed the absolute time difference in onset and offset of correctly predicted saccades and of saccades labeled by the human experts. This measure reflects how well an algorithm agrees with the human coder in terms of saccade start and end.

In all three datasets, U’n’Eye reached high similarity to the human coder (Fig. 4A, B, blue) and outperformed the EM algorithm (Fig. 4A, B, dark and light pink; Table 2). This was the case independent of how the EM *λ* parameter was optimized. U’n’Eye also detected saccade onset and offset in high agreement with the human labels. On average, saccade onset differences to human labels were smaller than 3 ms, and saccade offset differences were smaller than 5 ms. Saccade onset and offset labels by the EM algorithm deviated more strongly from the human-labeled saccades (Fig. 4C, D, blue vs. pink; Table 2). This indicates that U’n’Eye saccade predictions were more human-like than those of the threshold-based EM algorithm.

**Figure 4:**
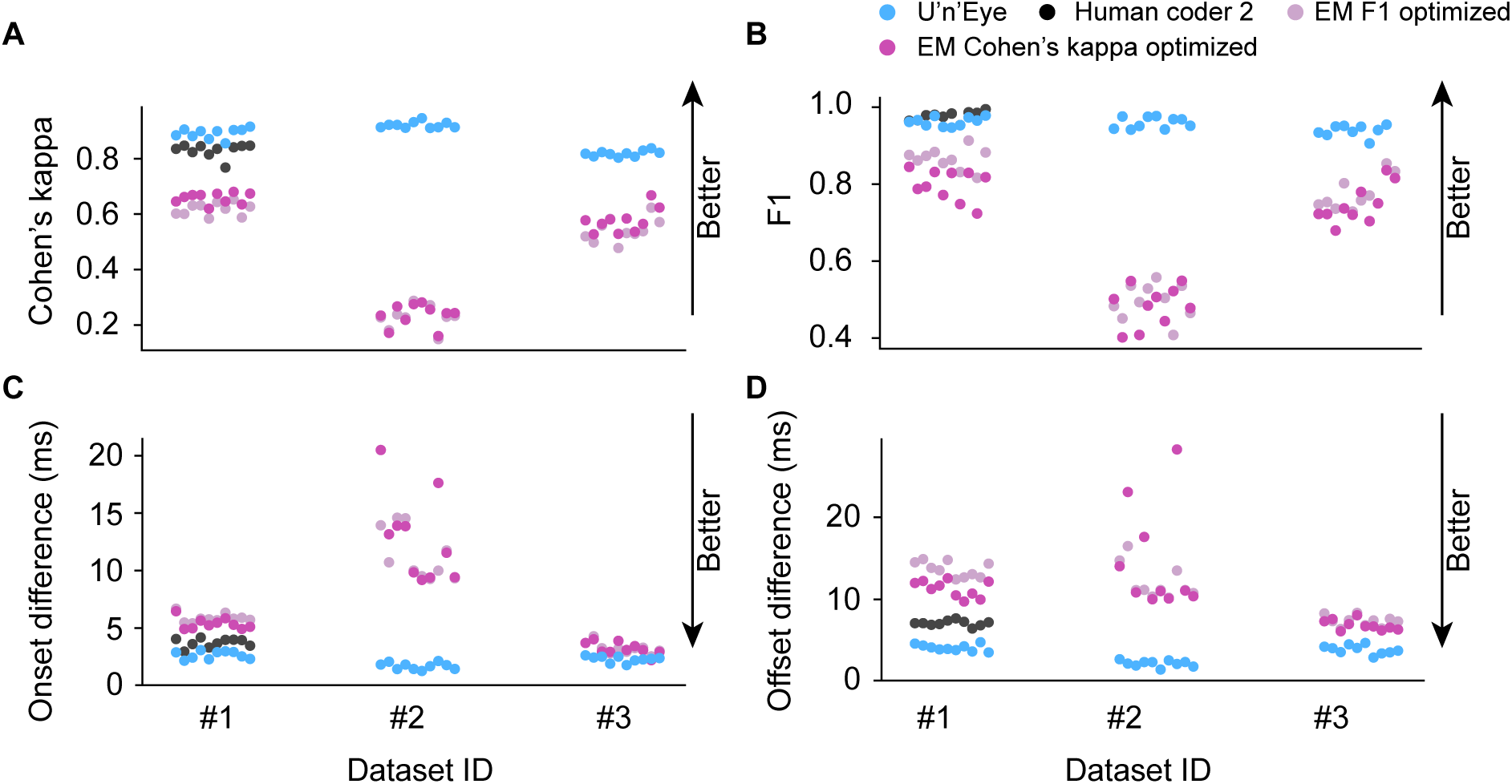
High performance of U’n’Eye. Each panel shows results from one performance metric described in the text, and on each of the three datasets. For each metric, we show results from 10 different cross-validation runs on the same 10% of data. (A) Cohen’s kappa measuring sample to sample agreement between two predictors. The first predictor was always a human coder. The second predictor (different colors) was either U’n’Eye (blue), a second human coder (black), the EM algorithm optimized to maximize Cohen’s kappa scores (dark pink), or the EM algorithm optimized for F1 (light pink). (B) F1 score summarizing precision and recall performance. (C) Average absolute difference in the timing of saccade onset times. (D) Same as C but for saccade offset times. In all cases, our network demonstrated superior performance.

**Table 2:** Comparison of U’n’Eye performance to the EM algorithm on the three datasets. EM (kappa) is the EM algorithm optimized to maximize the Cohen’s kappa metric. EM (F1) is the EM algorithm optimized to maximize the F1 metric. In bold are the best performances for each dataset. In all cases, U’n’Eye outperformed the EM algorithm. Values report mean and standard deviation across cross-validations.

| Dataset | Algorithm | Cohen's kappa | F1 | $\Delta$ Onset (ms) | $\Delta$ Offset (ms) |
| --- | --- | --- | --- | --- | --- |
| #1 | U'n'Eye | **0.89** ± 0.02 | **0.96** ± 0.01 | **2.66** ± 0.34 | **4.11** ± 0.41 |
| | EM (kappa) | 0.66 ± 0.02 | 0.80 ± 0.04 | 5.39 ± 0.49 | 11.28 ± 1.00 |
| | EM (F1) | 0.62 ± 0.02 | 0.87 ± 0.03 | 5.87 ± 0.38 | 13.68 ± 0.94 |
| #2 | U'n'Eye | **0.92** ± 0.01 | **0.96** ± 0.01 | **1.70** ± 0.29 | **2.19** ± 0.37 |
| | EM (kappa) | 0.23 ± 0.04 | 0.48 ± 0.05 | 12.85 ± 3.82 | 14.64 ± 6.38 |
| | EM (F1) | 0.23 ± 0.03 | 0.50 ± 0.05 | 11.37 ± 2.19 | 12.06 ± 2.12 |
| #3 | U'n'Eye | **0.82** ± 0.01 | **0.94** ± 0.01 | **2.30** ± 0.27 | **3.88** ± 0.54 |
| | EM (kappa) | 0.58 ± 0.04 | 0.75 ± 0.05 | 3.23 ± 0.55 | 6.86 ± 0.63 |
| | EM (F1) | 0.54 ± 0.04 | 0.77 ± 0.04 | 3.24 ± 0.49 | 7.35 ± 0.66 |

In the more challenging dataset 2, in which saccades occurred during smooth pursuit eye movements, U’n’Eye substantially outperformed the EM algorithm. Here, saccade velocity was close to the instantaneous velocity of the ongoing smooth pursuit movements. In fact, the minimum saccade peak velocity in this dataset was smaller than the median instantaneous velocity during pursuit (Table 1). This explains why the threshold-based EM algorithm failed to detect a large fraction of saccades in dataset 2, whereas U’n’Eye performed the best on this dataset when compared to the other datasets. This was because the network architecture utilizes a substantial time window (Fig. 2), which allows it to infer changes in the state of the eye even if the instantaneous velocity is low compared to the surrounding eye trace.

We next addressed the question whether U’n’Eye can achieve a similar level of inter-human agreement when multiple human experts analyze the same data. For this, we used dataset 1, because, among the three datasets, it contained saccades with the widest range of amplitudes (from as small as 0.02*◦* up to a size of 11*◦*; see Table 1 for a reason why saccades as small as 0.02*◦* were possible). We could thus assess inter-rater agreement for a broad range of saccades. Dataset 1 was labeled by a second independent human coder (Fig. 3, upper panel; Fig. 4, dataset 1). Coder 1 estimated saccade timing based on a combination of the raw eye traces and the smoothed radial velocity, whereas coder 2 used the raw eye traces only. We trained independent networks either with labels from coder 1 or coder 2 (Net 1 and Net 2, respectively), and we tested the networks’ performance on the 10 test sets from the 10-fold cross validation routine described above, both against ground truth labels from coder 1 or coder 2. U’n’Eye‘s saccade labels were as similar to both human coders as the human labels were to each other (Table 3). In terms of the F1 score, the inter-human agreement was not significantly different from the network-human agreement (Table 4). Interestingly, Net 1 showed higher similarity scores than coder 2 when both were compared to labels of coder 1 in the test sets, and vice versa for Net 2 and coder 2, reflected by larger Cohen’s kappa scores and smaller onset and offset differences (Table 4, all *p* < 5 · 10^−5^ after Bonferroni correction for multiple comparisons, Student’s paired samples t-test for Cohen’s Kappa and F1 scores, and independent samples t-test for on- and offset differences). This indicates that U’n’Eye‘s saccade estimation surpasses inter-rater consistency.

**Table 3:** Inter-rater comparison. The first row shows the similarity measures between labels from two human experts (coder 1 and 2). Net 1 was trained on labels from coder 1, and net 2 was trained on labels from coder 2. In bold are comparisons leading to best performances. Values report mean and standard deviation across cross-validations. Inter-coder agreement was evaluated on the 10 test samples from cross-validation.

| | Cohen's kappa | F1 | $\Delta$ Onset (ms) | $\Delta$ Offset (ms) |
| --- | --- | --- | --- | --- |
| Coder 1 vs Coder 2 | $0.83 \pm 0.02$ | **$0.98 \pm 0.01$** | $3.72 \pm 0.39$ | $7.10 \pm 0.34$ |
| Net 1 vs Coder 1 | **$0.89 \pm 0.02$** | $0.96 \pm 0.01$ | **$2.65 \pm 0.34$** | **$4.11 \pm 0.41$** |
| Net 2 vs. Coder 2 | **$0.89 \pm 0.01$** | $0.96 \pm 0.01$ | **$2.00 \pm 0.11$** | **$4.81 \pm 0.33$** |
| Net 2 vs. Coder 1 | $0.85 \pm 0.01$ | $0.96 \pm 0.01$ | $3.34 \pm 0.34$ | $5.58 \pm 0.33$ |
| Net 1 vs. Coder 2 | $0.86 \pm 0.01$ | $0.96 \pm 0.01$ | $2.82 \pm 0.32$ | $6.57 \pm 0.53$ |

**Table 4:** Statistical tests in inter-rater comparison. Net 1 (N1) was trained on labels from coder 1 (C1), and net 2 (N2) was trained on labels from coder 2 (C2). All p-values were Bonferroni corrected for multiple comparisons.

| Metric | Comparison | Test | t-value | p-value |
| --- | --- | --- | --- | --- |
| Kappa to C1 | N1 vs C2 | paired t-test | 18.38 | $2.98 \cdot 10^{-7}$ |
| Kappa rel. to C2 | N2 vs C1 | paired t-test | 10.88 | $3.69 \cdot 10^{-10}$ |
| F1 rel. to C1 | N1 vs C2 | paired t-test | -3.52 | $5.08 \cdot 10^{-2}$ |
| F1 rel. to C2 | N2 vs C1 | paired t-test | -3.7 | $5.19 \cdot 10^{-2}$ |
| Onset distance rel. to C1 | N1 vs C2 | indep. t-test | -6.6 | $2.98 \cdot 10^{-5}$ |
| Onset distance rel. to C2 | N2 vs C1 | indep. t-test | -13.6 | $5.26 \cdot 10^{-10}$ |
| Offset distance rel. to C1 | N1 vs C2 | indep. t-test | -17.9 | $5.28 \cdot 10^{-12}$ |
| Offset distance rel. to C2 | N2 vs C1 | indep. t-test | -15.3 | $7.33 \cdot 10^{-11}$ |

### U’n’Eye misses only a small fraction of microsaccades

We then analyzed the patterns of agreement and disagreement between U’n’Eye and human labeling. For true positive saccades, the two dimensional histogram of detected movements reflected the typical main sequence relationship between peak velocity and amplitude of saccades (Fig. 5,A,D,G)^40^. A few false positives were present within the range of the main sequence, suggesting that the human coder forgot to label some saccades (for example, see the movement in the inset in Fig. 5B). Concerning the rare false negatives that occurred, some of them had fairly large amplitudes (beyond eye tracker noise). Closer inspection revealed that there were pairs of successive saccades that had very short inter-saccadic intervals. The network lumped them into one movement, whereas the human coders separated them. Most remaining disagreements between the human and the network were associated with the smallest microsaccades, closest to eye tracker noise levels.

**Figure 5:**
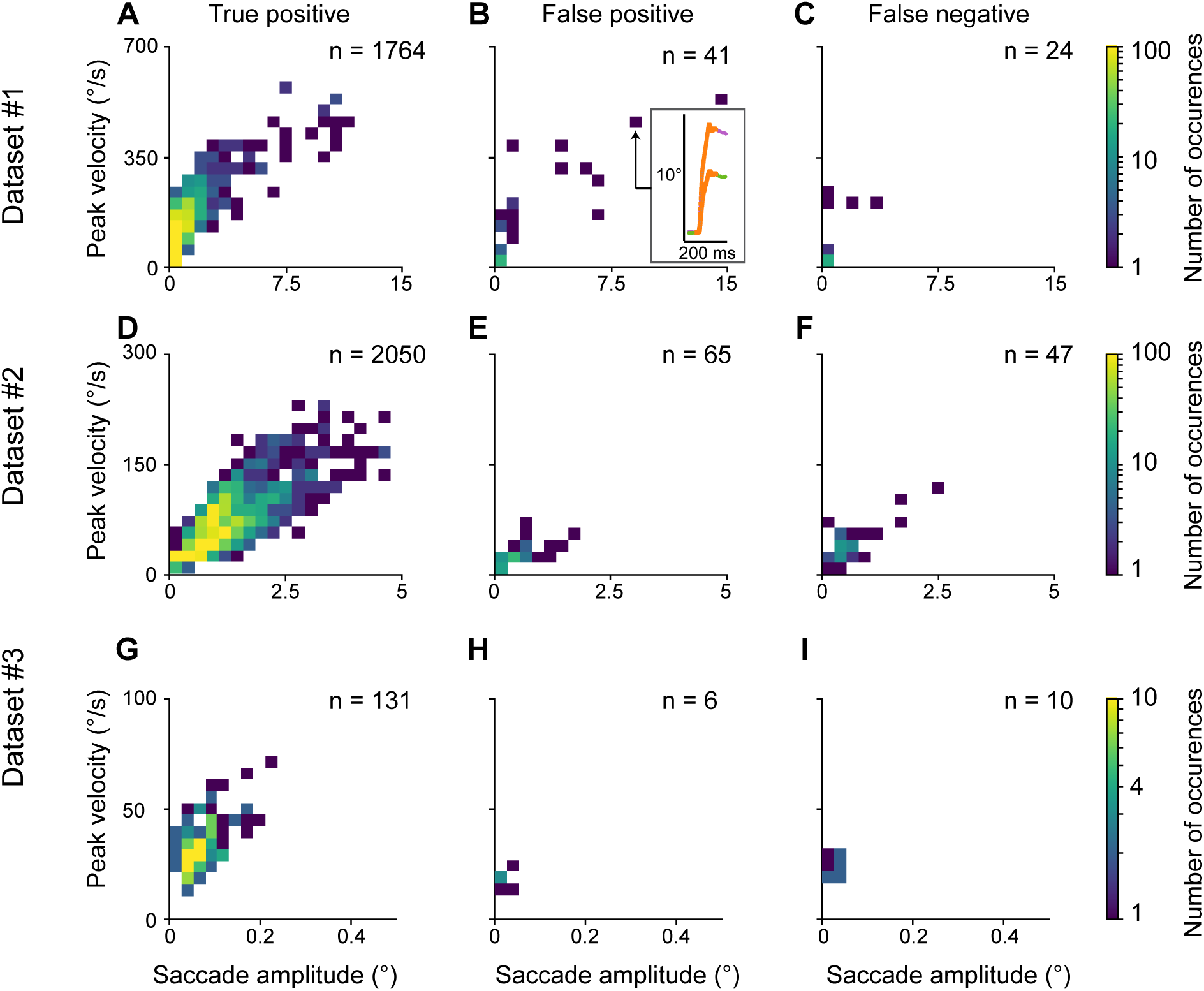
Location on the main sequence^40^ of detected and undetected saccades. Left panels show saccades that were detected both by a human expert and U’n’Eye. The detected saccades expectedly followed the main sequence relationship between peak velocity and movement amplitude. Middle panels show saccades that were detected only by U’n’Eye. Most saccades were small and close to the eye tracker noise, likely being cautiously unlabeled by human coders. In the inset, a large saccade was detected by U’n’Eye but not by the human coder, suggesting a possible lapse by the latter. Right panels show saccades missed by U’n’Eye. Most of these were very small.

### U’n’Eye: new state-of-the-art eye movement classifier

In order to compare our algorithm to state-of-the-art methods for eye movement classification, we evaluated its performance on a benchmark dataset^31^, which has previously been used for the comparison of 12 eye movement classifiers^29, 32^. The dataset comprises 500 Hz eye tracking recordings from humans watching videos, images, or moving dots, and it contains human labels for fixations, smooth pursuits, saccades, PSO (Fig. 6A), and blinks. We therefore used U’n’Eye as a multi-class classifier to predict saccades, PSOs, and blinks (Fig. 6B). Fixations and smooth-pursuit eye movements were both assigned to the fixation class. U’n’Eye output a predictive probability for each class (Fig. 6D), with the prediction value corresponding to the class that maximized this predictive probability (Fig. 6C). We trained U’n’Eye on one part of the data and evaluated its performance on the test trials listed in Andersson et al. (their Table 11^29^). U’n’Eye outperformed the state-of-the-art classifiers for saccades and PSOs (Table 5). Moreover, U’n’Eye’s performance lied within the range of the inter-coder agreement of the two human experts who labeled the dataset (Table 5). This result indicates that U’n’Eye is very well suited for multi-class eye movement classification.

**Figure 6:**
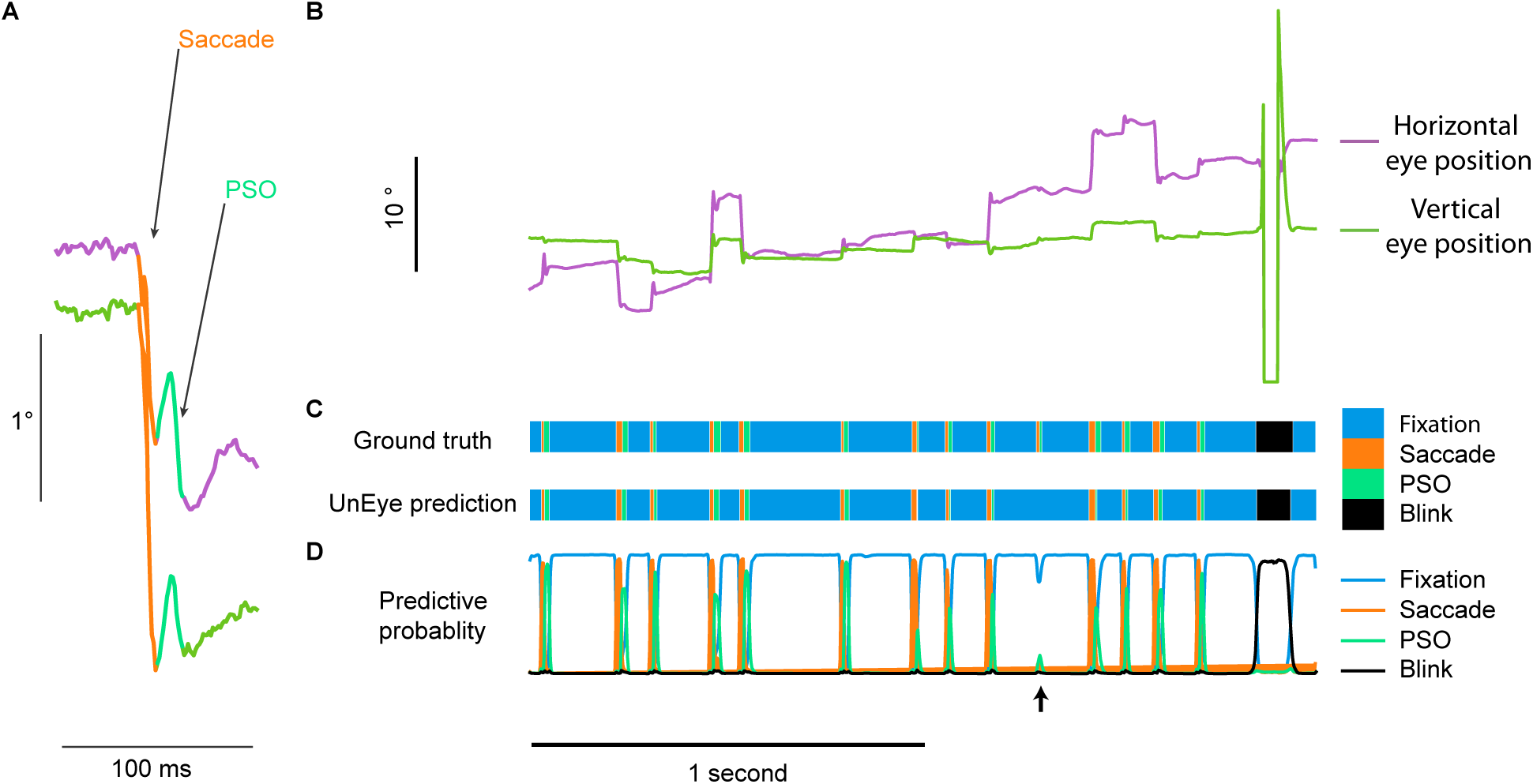
Multiclass labeling by U’n’Eye. (A) An example saccade showing substantial post-saccadic oscillation (PSO) from the data in ^31^. (B) An example full trace from the same dataset showing sequences of saccades, PSO’s, and blinks. (C) For the trace in B, ground truth labels are shown, in addition to labels by U’n’Eye. The latter successfully classified all ground truth labels, except for one instance marked by a black vertical arrow. (D) Nonetheless, the predictive probabilility of the network still showed a transient for the missed microsaccade (black arrow), suggesting that additional post-processing may be used to improve the performance of U’n’Eye even more.

**Table 5:** Performance of U’n’Eye compared to state-of-the-art algorithms. Bold face indicates highest values of an algorithm for each class. NSLR-HMM and LNS values were taken from the respective publication (NSLR-HMM^32^, LNS^29^). For U’n’Eye, values are the median across 20 independent networks.

|  | saccades | PSO | blinks |
| --- | --- | --- | --- |
| U'n'Eye | **0.89** | **0.71** | 0.83 |
| Coder MN | 0.90 | 0.73 | 0.91 |
| NSLR-HMM<sup>32</sup> | 0.82 | 0.42 | - |
| LNS<sup>29</sup> | 0.81 | 0.64 | - |

### U’n’Eye performance is robust to missing labels

In machine learning, the quality of the training data is crucial for the performance of a classifier, since the latter directly learns from the human ground truth labels. Human labeling, however, is prone to mistakes and lapses: saccades might be missed by the human coder, leading to noisy labels. We therefore assessed how U’n’Eye’s performance was influenced by noise-corrupted labels. We trained the network on simulated data (Fig. 1D) for which we knew the ground truth. We then artificially removed a fixed fraction of saccade labels. We also evaluated the network’s performance when trained on real data with missing labels. U’n’Eye was robust to the presence of noisy labels (Fig. 7). The removal of 20% of saccade labels during training led to a decrease of only 0.037 in Cohens kappa and 0.028 in F1. Moreover, there was only 0.59 ms and 1.01 ms differences in onset and offset estimation, respectively (Fig. 7A, B). This indicates that U’n’Eye can achieve good performance even if the human coder misses some saccades in the training set.

**Figure 7:**
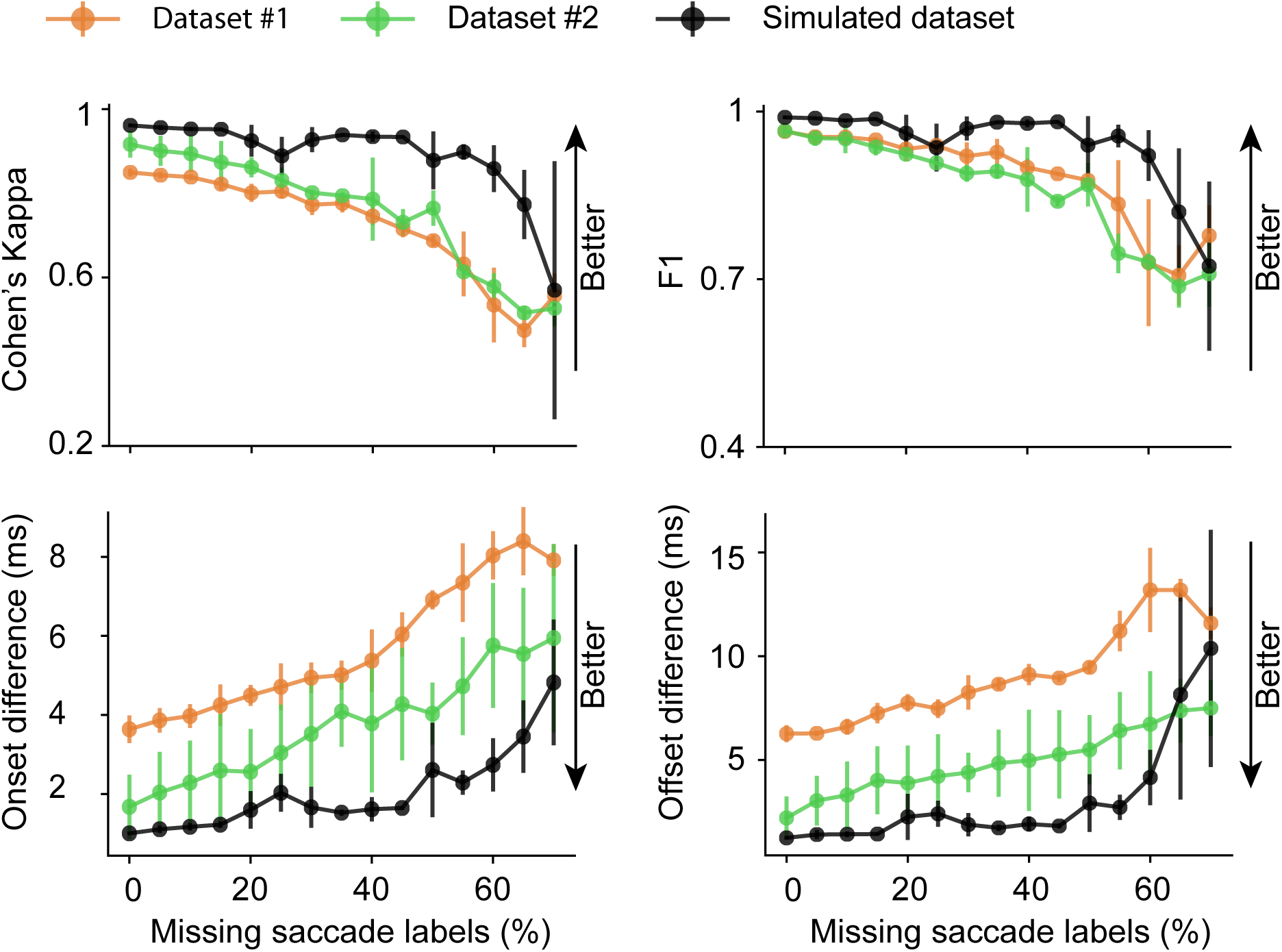
Classification accuracy of networks trained with random missing labels. We used the same performance metrics of Fig. 4, and we computed these metrics on the outputs of networks trained on a simulated dataset (Fig. 1D) as well as on datasets 1 and 2. The simulated dataset generated ground truth eye traces (with noise added later; Methods) using a model adapted from ^41^. For all datasets, we systematically removed fractions of labels from the training set to evaluate degradations in U’n’Eye performance. Classification accuracy measured in terms of Cohen’s Kappa (top row, left) and F1 score (top row,right) decreased with increasing label noise. The bottom row shows performance in estimating saccade onset and offset times. Note that performance in all metrics remained within a high range even with high noise fractions. The data show mean standard deviation for 5 independent training and test procedures.

### Eye movement representation becomes disentangled along network layers

We finally had a closer look at how the network achieves the separation of two eye states (e.g. fixations and saccades; Fig. 8A). In the velocity domain, saccades and fixations can show highly overlapping distributions (Fig. 8B). This explains why velocity threshold-based algorithms can fail to distinguish fixations from saccades (Fig. 4). Here, we showed that U’n’Eye can differentiate between fixations and saccades with high accuracy (Fig. 4). The classification was based on the output layer of the network. To illustrate how this decision arises throughout the hidden layers, we performed principal component analysis (PCA) on the features of each convolutional layer. The fraction of explained variance by the first two principal components (PCs) reflects the U-shaped architecture of the network (Fig. 8C): in the middle layers, information is distributed across more components than in early and late layers. We projected the hidden layer activations onto the PC space and la-beled time bins according to their ground truth labels (fixation or saccade, Fig. 8D). We observed in higher layers that the two classes were better separated (Fig. 8D). Finally, in the output layer, fixations and saccades became linearly separable (Fig. 8E). Thus, through training, the network effectively learns to extract relevant features and to project those onto a plane where the two eye movement classes are linearly separable.

**Figure 8:**
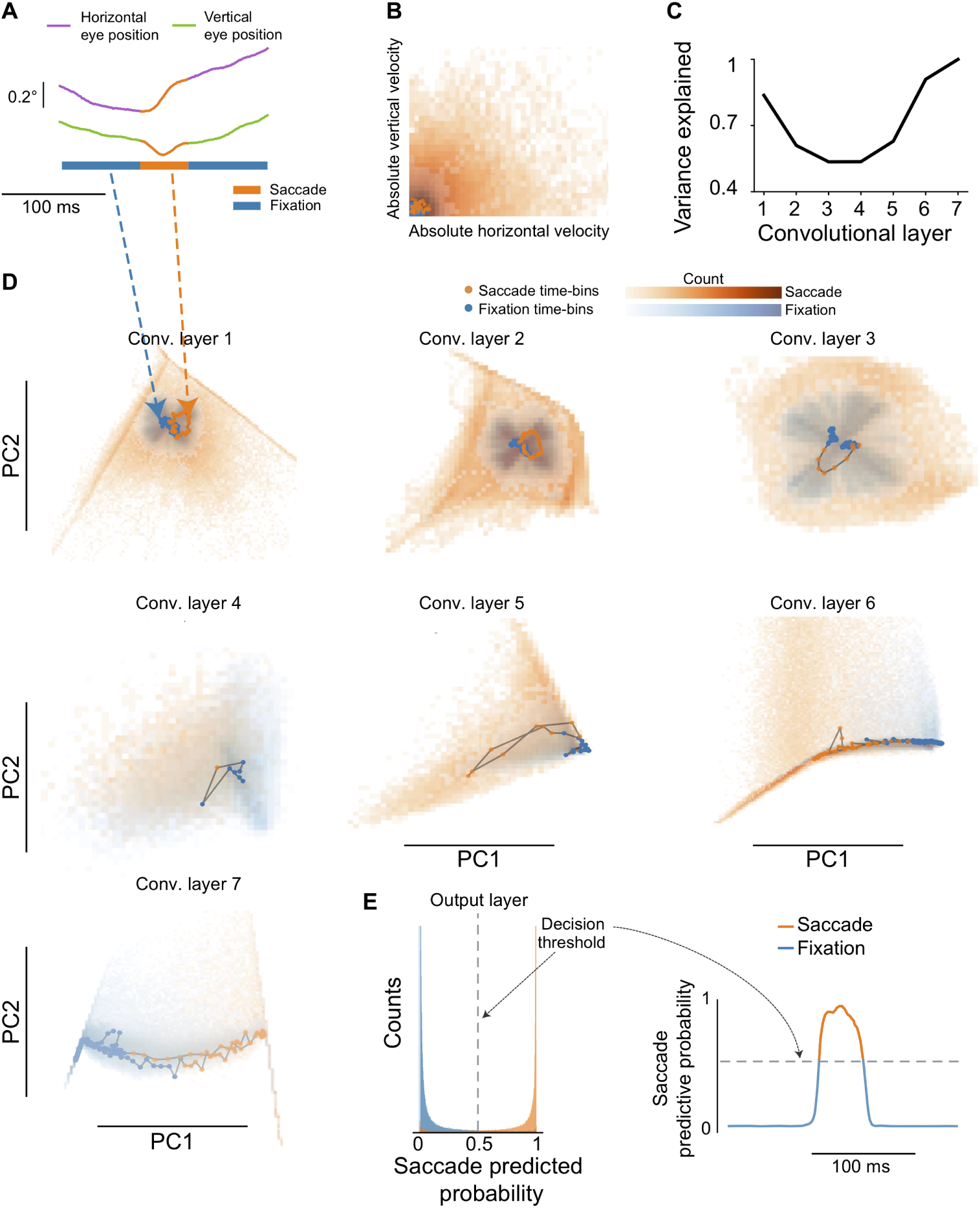
Disentanglement of fixations and saccades throughout the network. (A) Example eye trace with a microsaccade. B) Distribution of dataset 2 in the velocity domain. Fixations and saccades (shown in bluish and orangish colors, respectively) showed overlapping distributions. (C) Fraction of explained variance by the two first principal components (PCs) of the network’s convolutional layers. There was a reduction in the middle layers followed by a peak at the final seventh layer. (D) Projection of hidden layer activations by eye traces of dataset 2 onto the first two principal components. Fixation and saccade classes became better separated throughout the hidden layers. (B – D) Dots indicate the time points of the example eye trace in A, and the rest of the background data show the entire dataset time samples. (E) The probability output allowed for a linear separation of the two classes. Time points with a saccade predictive probability above 0.5 were classified as a saccade.

## Discussion

In this study, we presented U’n’Eye, a convolutional neural network for eye movement classification. We demonstrated that U’n’Eye achieved human-level performance in the detection of saccades and microsaccades. In addition, the network was able to predict other classes of eye movements, which we exemplified with the detection of blinks and PSOs in a benchmark dataset.

Furthermore, we showed that U’n’Eye achieved excellent performance both when trained on a single type of data with labels from one coder and when trained on different datasets with labels from two coders. While datasets 1 and 3 used in this study contained data with only one type of visual task and labels from one coder each, dataset 2 was composed of two different pursuit tasks and contained labels from two different human coders. The dataset by Andersson et al. also contained greatly varying types of saccades and other eye movements. Still, U’n’Eye achieved good performance when trained and tested on this dataset. Note that the network might fail to detect eye movements when tested on data that show a different distribution than the data it was trained on. We therefore recommend to either train a network with a large variety of data or to train separate specialized networks for each task.

U’n’Eye is publicly available and provides a user friendly interface as well as a web service in which users can upload their data and receive classification outputs. No parameter tuning is needed even for training (e.g. learning rate, and so on) since the standard settings were found to work well across datasets. Instead, an experimenter just needs to provide a few hundred seconds of labeled data to train the network once. Even if some labels are missing in the training data, U’n’Eye can still reach high performance. We recommend, however, to use only carefully annotated data for training, as this will improve results.

Of the few algorithm that are capable of detecting saccades as well as PSO^31, 32, 42^, U’n’Eye achieves highest performance. Note that Zemblys et al. ^42^ reported higher absolute Cohen’s kappa values for saccade and PSO detection, but obtained these on cleaner data than the benchmark dataset from Andersson et al. ^29^. As their dataset was not available to us, we were not able to assess U’n’Eye‘s performance on the same data.

Recently, a Bayesian approach for the detection of microsaccades based on a generative model has been proposed^43^. Inherently, Bayesian methods provide estimates of uncertainty, in addition to estimates of the quantity of interest. Indeed, it is an interesting future perspective to combine U’n’Eye with Bayesian Deep Learning techniques to provide uncertainty estimates for the detected eye movements^44^.

Future work will include combining datasets with different characteristics, such as different sampling frequencies, in order to obtain a network that can generalize on a large range of data. Such a network could be used by a large part of the scientific community, which would allow for reproducibility of scientific results. We recommend that anyone who uses our algorithm to publish the weights of the trained network so that eye movement detection can be reproduced. For our own trained networks, all weights have been published online (https://github.com/berenslab/uneye) along with the code of the network. This has the advantage that users with similar data characteristics to one of our three datasets (e.g. microsaccades during fixation with a video-based eye tracker as in Dataset 3) can directly use our weights from the proper dataset without having to retrain their own network. We will also make all three datasets publicly available, facilitating the further development for eye movement detection algorithms.

Of course, it should be noted that some prediction errors may still occur with U’n’Eye. However, such errors fall within the range of inter-rater variability across humans anyway. Also, even when U’n’Eye does make mistakes, the predictive probability that it outputs can be used to retrieve missed saccades (e.g. see the black arrow in figure Fig. 6). Therefore, post-processing algorithms may be used to further optimize the output of U’n’Eye.

Finally, U’n’Eye’s capacity to learn non-linear relationships between an eye trace and some annotated labels opens new horizons in neuroscience: the network could be used to understand the properties of neural activities related in a complex manner to eye movements. For example, the disentanglement in later layers (Fig. 8) could be used to quantitatively analyze the activity patterns of pre-motor neurons in the brainstem, which themselves ultimately transform brain processing into individual ocular muscle innervations. Furthermore, U’n’Eye could be turned into a generative model for eye movements, as was shown for neural networks that are used for image classification ^45^. The information about eye movements that is contained in the network architecture might in the future be used to identify variations in eye movement characteristics that could hint at underlying pathologies.

## Methods

### Datasets

All experiments used for collecting the datasets were approved by ethics committees at Tuebingen University. Human subjects provided informed, written consent in accordance with the Declaration of Helsinki. Monkey experiments were approved by the regional governmental offices of the city of Tübingen.

Dataset 1 was collected from human subjects using the Eyelink 1000 video-based eye tracker (SR Research, Ltd) sampling eye position at 1 kHz. The dataset contains mostly microsaccades and small-amplitude memory-guided saccades.

Dataset 2 was collected from three male, rhesus macaque monkeys implanted with scleral search coils (in one eye for each of the monkeys). The dataset contains catch-up saccades generated during smooth pursuit. Eye position was again sampled at 1 khz. For the trials containing smooth pursuit of sinusoidal target motion trajectories in this dataset, the data were obtained from the experiments described in ^46, 47^.

Dataset 3 was collected from a single male macaque monkey using the Eyelink 1000 video-based system sampling eye position at 500 Hz. The dataset contains microsaccades generated during fixation. The data was obtained from experiments described in ^48^.

In all datasets, we manually detected saccades using a custom-made GUI in Matlab. The GUI displayed horizontal and vertical eye position traces, as well as filtered radial eye velocity. The GUI internally estimated saccade onset and end times using a combination of velocity and acceleration thresholds^49^. The user then manually interacted with the GUI to delete false alarms, correct false negatives, and adjust estimation of onset and offset timing.

### Simulated saccades

To test the performance of our network on noisy labeled data, we designed artificial eye traces for which we knew the ground truth. Saccades ranging from 0.5 to 60*◦* were simulated using an adaptation of a model for saccade waveforms^41^. The model is a sum of a soft ramp function, which follows the relationship between amplitude and peak velocity observed in real saccades^41^. As it is originally one dimensional, we adapted it so that it generates two dimensional trajectories. Saccade generation in time was made to follow a Poisson process with *λ* equal to 3 saccades per seconds. Simulated blinks were also added by inducing sharp transients in the eye traces. Finally, a Gaussian white noise with a standard deviation of 0.02*◦* was added to the trace. Then, as described in Results, we trained U’n’Eye under a variety of conditions in which we intentionally removed a subset of saccade labels during training, in order to explore robustness to missing labels (Fig. 7).

### U’n’Eye: our convolutional neural network

The architecture of the convolutional neural network (CNN) was inspired by U-Net, a CNN first used for image segmentation^33^. Here we modified U-Net to meet the requirements of an eye movement classifier. The network was built of seven convolutional layers with kernel size 5, each followed by a linear-rectifying unit (ReLU) and a BatchNorm layer, both described in detail in Results. Batches consisted of samples of the same duration. The input to the network was eye velocity, as a matrix of dimension N x T x 2, where N is the batch-size, T the number of time-points and 2 the number of coordinates (horizontal and vertical eye velocity). The number of input time-points could be variable but had to be a multiple of 25 bins due to the max pooling operations. The output of the network was a matrix of dimension N x K x T, where K was the user-defined number of classes. For example, we could have a “saccade” and “fixation” class in the networks of Fig. 3 and we could also add other classes like “PSO” in the network of Fig. 6.

We applied a softmax^38^ activation function to the output of the last convolutional layer x:

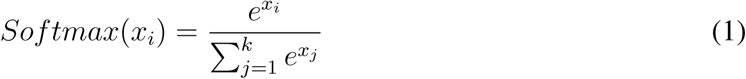

where *x_i_* is the layer corresponding to class i. Thus, the network output y represented the sample-by-sample conditional probability of each class (e.g. “fixation” or “saccade”) given the eye-velocity x and the network weights w:

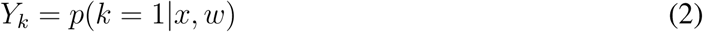

The final prediction of the algorithm represented the class that maximized this conditional proba-bility:

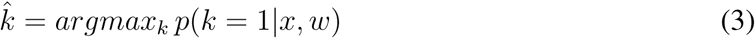

We chose the kernel sizes of the convolutional and max pooling operations in a way to capture a relevant signal range around each time-point. Based on the given kernel sizes of the network, it can be shown that the prediction of one time-bin is influenced by the preceding and following 89 time-bins of the velocity signal (2B, red color).

### Network training

We trained the network with mini-batches whose size depended on the total number of training samples. We performed 10 training iterations in each epoch. Over-fitting on the training set was prevented by computing the loss on a validation set and stopping training when the validation loss increased for three successive epochs. We used a multi-class error function which, for two classes, equals the cross entropy loss. Weight-regularization was done with L2-penalty^38^, which corresponds to a Gaussian prior with zero mean over the network weights. The optimal parameter *λ* was determined to be 0.01. The loss function was thus defined as:

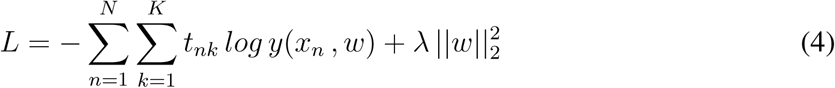

where N is the number of time points and K the number of classes. The ground truth label *t_nk_* equals 1 if the time point n belongs to class k. Gradient computation was done with PyTorch autograd method.

We use Adam optimizer ^50^ with an initial learning rate of 0.001. Adam is a stochastic gradient-based optimizer that uses adaptive learning rates for different weights of the network. An additional step-decay by a factor of 2 was applied to the current learning rates when the loss on the validation set increased during one epoch.

### Postprocessing

In the case of binary prediction into the classes fixation and saccade, we provided the possibility to define thresholds for minimum saccade duration and minimum saccade distance. If thresholds were given, saccades closer than the minimum distance were merged and saccades shorter than the minimum duration were removed. We obtained the results reported here with a minimum saccade distance threshold of 10 ms for dataset 1 and 3 and 5 ms for dataset 2, because we previously observed that some saccades occurred very close in time in this dataset. For datasets 1–3, we used a minimum saccade duration threshold of 6 ms. The same thresholds were used for the algorithm by Engbert & Mergenthaler^30^.

### Data augmentation

U’n’Eye performs better with a bigger training set. However, we aimed to reduce the amount of saccades that a user should provide to train U’n’Eye. In this study, to increase the number of training samples, the input eye positions were rotated and added to the original training samples:

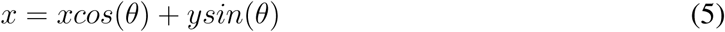

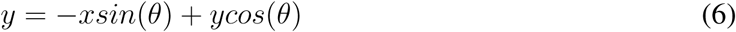

where x and y are the horizontal and vertical eye positions. We used *θ*=(1/4*π*,3/4*π,*5/4*π*,7/4*π*). Thus, we could increase by five fold the size of our training set without causing over-fitting.

### Performance measures

To evaluate the eye movement detection performance of our network, we used the following metrics: CohenâĂŹs kappa, F1 score, and onset and offset time differences.

CohenâĂŹs kappa is a sample-based statistic. It reflects how much two coders agree on the class that each time-bin belongs to, while controlling for chance agreement of the two coders. It is given by:

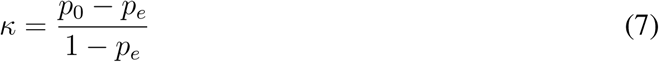

where *p*_0_ is the proportion of time-bins for which two coders agree and *p_e_* is the proportion of time-bins for which agreement can be expected by chance.

For a binary classification of fixation versus saccades, the CohenâĂŹs kappa value *p_e_* is given by

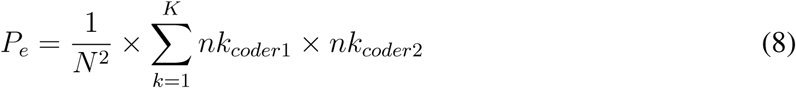

where *nk_coderX_* is the number of time-bins coderX assigned to class k.

The F1 score is a measure of classification accuracy that combines precision and recall of a predictor. Precision is defined as the proportion of correctly classified saccades over all predicted saccades. Recall is defined as the proportion of correctly classified saccades over all saccades in the ground truth. The F1 score is the harmonic mean of these two measures. It is given by:

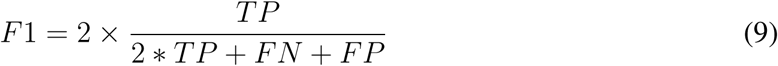

where TP is the number of true positives, FN the number of false negatives, and FP the number of false negatives. For all true positive saccades, we compared saccade timing between the ground truth and prediction by calculating the absolute time differences between true and predicted saccade onsets and offsets.

### Evaluation on a benchmark dataset

We evaluated U’n’Eye performance on a benchmark dataset by Andersson et al.^29^. This dataset comprises 500 Hz eye-tracking data from humans looking at images, movies, or moving dots. It contains human labels for the events fixations, smooth pursuits, saccades, PSO and blinks. Events that the human experts did not assign to any of these classes were labelled as “others”. For some trials, the dataset contained labels from two different human coders. For other trials, only one label was available. We trained 20 independent networks with different random initializations on the data with labels from one human coder (coder RA). Performance was then tested on the trials with labels from two coders, which makes our result comparable with previously reported results^32^. Note that we were not able to reproduce the inter-rater measures reported by Andersson et al.^29^ in line with the results of Pekkanen and Lappi^32^. For comparability with the NSLR-HMM algorithm^32^, we excluded the event labels “blinks” and “other” for the calculation of the saccade and PSO Cohen’s kappa scores. The Cohen’s Kappa scores for blinks were calculated excluding the label “other”. However, the performance on this class was not compared to other algorithms since it was not reported.

### Compute time

The computation times of our algorithm reported here were achieved on a personal computer with a 3.1 GHz Intel Core i5 processor at 16 GB RAM running on Mac OS X 10.11.6.

### Code and data availability

All code is available from https://github.com/berenslab/uneye. Data as well as a web service will be available upon publication.

## Acknowledgements

We thank Konstantin-Friedrich Willeke and Antimo Buonocore for providing help with labeling saccade data and Murat Ayhan for input on DNNs. This work was funded by the German Ministry of Education and Research (FKZ 01GQ1601) and the German Research Foundation (EXC307, SFB 1233, BE5601/4-1).

## Author contributions

MB, JB, ZH and PB designed the project; MB and JB developed the algorithm; MB implemented the algorithm; JB simulated, acquired and labeled data; MB and JB analyzed the data; HN provided data; ZH and PB supervised the project; MB, JB, ZH and PB wrote the paper.

